# Drug-Target Interaction Prediction with PIGLET

**DOI:** 10.64898/2026.02.18.706530

**Authors:** Kristy A. Carpenter, Russ B. Altman

## Abstract

Drug-target interaction (DTI) prediction is a key task for computed-aided drug development that has been widely approached by deep learning models. Despite extremely high reported performance, these models have yet to find widespread success in accelerating real-world drug discovery. In contrast with the most common approach of creating embeddings from one-dimensional or three-dimensional representations of the input drug and input target, we create a novel graph transformer method for DTI prediction that operates on a proteome-wide knowledge graph of binding pocket similarity, protein-protein interactions, drug similarity, and known binding relationships. We benchmark our method, named PIGLET, against existing DTI prediction models on the Human dataset. We assess performance with two different splitting strategies: the frequently-reported random split, and a novel, more rigorous drug-based split. All models perform similarly well on the random split, and PIGLET outperforms all models on the drug-based split. We highlight the utility of PIGLET through a real-world drug discovery case study.

## 1 Introduction

An oft-cited promise of machine learning for drug discovery is the ability to expeditiously conduct virtual screening [4]. Besides using machine learning methods to enhance physical docking calculations [21, 22] and molecular dynamics simulations [1, 2, 14, 31], two common formulations of the virtual screening prediction task are the drug-target affinity (DTA) task and the drug-target interaction (DTI) task. DTA is a regression task that requires predicting the binding affinity between a query target and query drug, usually in terms of inhibition constant (*K*_*i*_), dissociation constant (*K*_*d*_), or half-maximal inhibition concentration (*IC*_50_). DTI is a binary classification task that only requires predicting whether or not a binding interaction will occur between a query target and a query drug.

A large variety of methods, including deep learning methods, for DTI and DTA exist [8, 32]. One common approach for the DTI/DTA task is to learn a representation for the query drug and a representation for the query target, then fuse the two representations to predict an affinity or binary interaction. Representations can be one-dimensional. DeepDTA [44], the first DTA model, leveraged a convolutional neural network (CNN) architecture to predict affinity from featurized SMILES strings and amino acid sequences. Similarly, DeepConv-DTI [16] encoded amino acid sequences with a CNN and drug molecular fingerprints with a feedforward neural network and fused outputs with a feedforward neural network. When transformers emerged as a dominant deep learning architecture, DTA/DTA models transitioned away from CNNs. TransformerCPI [5] and AMMVF-DTI [35] both embedded amino acid sequences using word2vec and SMILES strings using RDKit before processing both with transformer architectures and predicting interaction probabilities. MolTrans [10] used transformers to learn representations of SMILES strings substructures and primary sequence substructures. DTI-BERT [41] leveraged a BERT model to extract features from amino acid sequences and drug molecular fingerprints and predict binary interactions. SMFF-DTA [37] used drug SMILES strings, drug Morgan fingerprints, target secondary structure, target primary sequence, and target physicochemical properties as input to a multiple attention block.

Three-dimensional drug and target representations are also very common. As with their one-dimensional counterparts, early deep learning approaches leveraged CNNs. Ragoza *et al*. [23], KDeep [11], and Pafnucy [25] all used 3D CNNs on protein-ligand complex structures to predict binding score or binding affinity. OnionNet [42] used a 2D CNN on structural features derived from concentric shells centered on the ligand interaction site. Later, graph convolutional networks (GCNs), in which drug atoms and target amino acids were represented as nodes in a graph, emerged as a popular approach. Torng and Altman [29] and 3DProtDTA [33] both leveraged GCNs on protein and ligand molecular graphs to score drug-target interactions. FragXsiteDTI [13] used GCNs to featurize binding pocket structures and drug substructures, then input these featurizations to transformer blocks to score the interaction and propose the most likely pocket and drug substructure interacting pair.

A less common approach for DTI/DTA is to model an interaction network of proteins and drugs. This approach explicitly leverages biological relationships and similarities to predict drug-target interactions, following a guilt-by-association principle. MSF-DTA [20] augments a 3D approach, in which the drug is represented as a molecular graph, by encoding targets using their protein-protein interactions and sequence similarity relationships. DHGT-DTI [36] uses a graph transformer over a heterogeneous network of drugs, targets, diseases, and side effects and predicts drug-target interactions through matrix factorization. TxGNN [9] uses a medical knowledge graph with biological process, protein, disease, phenotype, anatomy, molecular function, drug, cellular component, pathway, and exposure nodes, which is processed by heterogeneous graph convolutions. Drug-disease relationships are predicted through DistMult. While not an explicit DTI model, TxGNN embodies a similar approach for an adjacent task.

While powerful, network-based approaches to DTI/DTA are limited by the scope of existing biological knowledge. For example, MSF-DTA only used protein sequence similarity due to the lack of structural information, and the protein nodes in DHGT-DTI represent less than 20% of the human proteome due to the lack of functional information. In [3], we created a proteome-wide predicted pocket dataset based on both experimentally-determined and computationally-predicted protein structures. With this resource, we can identify similar binding pockets across the entire human proteome and take advantage of the principle that highly similar binding pockets are likely to bind similar ligands.

In this work, we create a heterogeneous knowledge graph comprising protein-protein interactions, protein-drug interactions, pocket-pocket similarity, and drug-drug similarity throughout the whole human proteome. To predict drug-target interactions based on this graph, we develop PIGLET (**P**roteome-wide **I**nteraction **G**raph **L**ink prediction by **E**mbedding with **T**ransformers), a graph neural network model comprising a graph transformer embedding trunk and a link prediction head. We show that PIGLET performs comparably to state-of-the-art DTI prediction models on the Human dataset [18] when randomly split. We create a more rigorous split of the Human dataset based on drug similarity and demonstrate the PIGLET outperforms other DTI prediction models on this split. Finally, we apply PIGLET to identify drugs with potential off-target binding to the mu opioid receptor. The PIGLET model is available at https://github.com/Helix-Research-Lab/PIGLET.

## 2 Methods

### 2.1 Datasets

We selected the Human dataset [18] to train, evaluate, and benchmark PIGLET due to its frequent use for benchmarking DTI models. This dataset consists of interactions between human proteins and drugs, contains high-quality negative examples, and is balanced between positive and negative examples.

In addition, we augmented our knowledge graph with DTI information from DrugBank [38]. We extract target information from all drug entries in DrugBank to obtain a set of positive DTI examples.

This data was not used for calculating loss or any evaluation, but rather to guide the inductive bias of PIGLET using message-passing.

### 2.2 Assembly of the proteome-wide interaction graph

We created a proteome-wide interaction graph upon which PIGLET conducted message passing and link prediction. This heterogeneous graph *G* can be represented as a set of vertices *V* and edges *E. G* is undirected and does not contain self-loops. *V* = {*V*_target_, *V*_drug_}, where each node 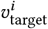 in *V*_target_ represents a different protein from the canonical human proteome and each node 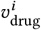 in *V*_drug_ represents a different drug. *E* = {*E*_PPI_, *E*_targetsim_, *E*_drugsim_, *E*_bind_, *E*_bindMP_}, where each edge 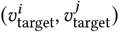 in *E*_PPI_ represents a protein-protein interaction between the *i*th target and the *j* th target, each edge 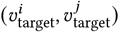 in *E*_targetsim_ represents high similarity between the binding pockets of the *i*th target and the *j* th target, each edge 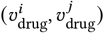 in *E*_drugsim_ represents high similarity between the *i*th drug and the *j* th drug, each edge 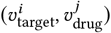 in *E*_bind_ represents a binding interaction between the *i*th target and the *j* th drug derived from the Human dataset, and each edge 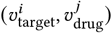 in *E*_bindMP_ represents a binding interaction between the *i*th target and the *j* th drug derived from the DrugBank dataset.

We used HOTPocket [3] to obtain predicted binding pockets for the canonical human proteome. Any target that was a member of the canonical human proteome as defined by UniProt [28] and that had at least one predicted pocket from HOTPocket or known pocket from the BioLiP2 database [40] was included as a node in *V*_target_. We used ESM2 [17] to obtain per-residue embeddings for each pocket, computed cosine similarities between the ESM2 embeddings of each residue in each pocket for all pairs of proteins, and defined any pair with a cosine similarity greater than 0.95 as a highly similar pair:

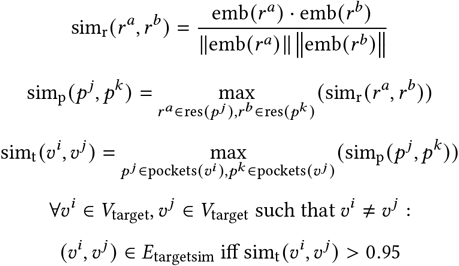

where emb (*r*) returns the ESM2 embedding for a residue *r*, res (*p*) returns all residues in a pocket *p*, and pockets (*v*) returns all HOTPocket pockets of all structures for a given target *v* which is also a node in *V* .

Highly similar target pairs, which were added as edges to *E*_targetsim_, comprised about 5% of all possible target pairs.

Edges in *E*_bind_ and *E*_bindMP_ were obtained from binding relationships in the Human dataset [18] and DrugBank dataset [38], respectively. We filtered both datasets to contain only targets represented in *V*_target_. All drugs were added to *V*_*drug*_ and all target-drug edges were added to *E*_bind_ and *E*_bindMP_. We used the ChemBERTa [6] embeddings of drugs as node features.

Drug-drug similarity was determined by the cosine similarities between ChemBERTa embeddings of drugs. Any pair of drugs with a cosine similarity greater than 0.8 was retained as a highly similar drug pair and added as an edge in *E*_drugsim_. Highly similar drug pairs comprised 4.98% of all possible drug pairs.

PPI information was obtained from the STRING database [26]. STRING contains PPIs from several different resource types: neighborhood, fusion, co-occurrence, experimental, database, and text mining. Each PPI in STRING is annotated with confidence scores for each of the resource types as well as a combined score. We filtered for PPIs between proteins represented in *V*_target_ with a combined score of at least 700. Of these, we retained PPIs. with an experimental score of at least 700 or a database score of at least 700. These PPI relationships were encoded in *G* as *E*_PPI_ edges.

### 2.3 PIGLET algorithm

The PIGLET architecture consists of two parts: an embedding trunk and a link prediction head. The embedding trunk conducts message passing over the proteome-wide interaction graph described above to learn embeddings for both target nodes and drug nodes. The link prediction head uses the node embeddings to predict whether or not an edge between a given drug-target pair exists. We train the whole model with binary cross-entropy loss and the Adam optimizer. A schematic of the PIGLET architecture and proteome-wide interaction graph is shown in Figure 1.

**Figure 1:**
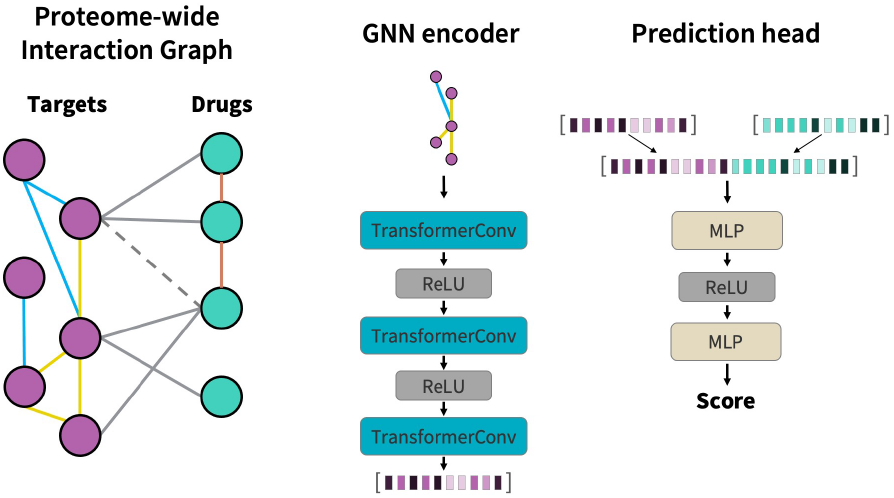
Schematic of PIGLET architecture. PIGLET is built on a proteome-wide interaction graph with nodes representing drugs and targets, and edges representing similarity and interaction relationships. The drug-target interaction prediction task is formulated as link prediction in this heterogeneous graph. The GNN encoder trunk of PIGLET learns node embeddings with transformer convolutions. The prediction head of PIGLET predict drug-target interaction scores by concatenating a drug node embedding and a target node embedding and passing it through a feedforward neural network.

#### 2.3.1 Embedding trunk

The embedding trunk is made up of 3 heterogeneous graph convolution layers, which learn different graph convolution parameters for each message-passing edge type in *E*. We introduce a virtual node connecting all drug nodes to facilitate information propagation. PIGLET uses the TransformerConv layer [24] to perform graph transformer message passing:

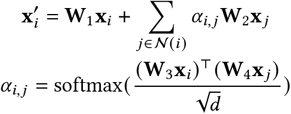

We use the ReLU activation function between TransformerConv layers.

#### 2.3.2 Link prediction head

The link prediction head receives two node embeddings as input. The embeddings are concatenated and passed to a two-layer feedforward neural network. We use the Leaky ReLU activation function between feedforward layers.

#### 2.3.3 Hyperparameter tuning

We performed hyperparameter tuning to select the batch size, learning rates, embedding dimension, graph convolution function, number of GNN embedding trunk layers, and number of link prediction head layers. We used the drug-based split with five-fold cross-validation (described below) for hyperparameter tuning. We did not use the testing set for any part of hyperparameter tuning.

### 2.4 Data splits

We used two different strategies to split the Human dataset into train, validation, and test sets. In both cases, we excluded all edges in *E*_bind_ from message passing; all training set binding edges were used for gradient descent. Edges in *E*_bindMP_ were the only binding edges used for message passing and were not used for gradient descent.

#### 2.4.1 Random split

Many existing DTI models report performance on random splits of the Human dataset [13, 30, 39, 43]. We created random 80:10:10 splits of the Human dataset to compare PIGLET to these models. Each model was evaluated on the same 20 random splits.

#### 2.4.2 Drug-based split

Random splits can give inflated estimates of model performance due to data leakage. In an effort to mitigate this, we created a split of the Human dataset based on drug similarity. A split strategy that holds out drugs simulates the real-world scenario of novel drugs being introduced, whereas a held-out target strategy is unneeded as structure predictions for the entire human proteome are currently available and new proteins will not be introduced on a human lifetime timescale.

We used RDKit to generate Morgan fingerprints of all drugs in the Human dataset. We performed a hierarchical clustering based on Tanimoto similarity between Morgan fingerprints. We cut the resulting dendrogram such that there were 100 clusters. We partitioned the clusters into five cross-validation folds (90% of data in total) and a held-out test set (10% of data) such that all binding interactions for a given drug belonged to that drug’s designated split. The held-out test set constituted 10% of the total data and contained 290 positive binding examples (48.33%) and 310 negative binding examples (51.67%). In total, the cross-validation folds constituted 90% of the total data and contained 2316 positive binding examples (43.70%) and 2984 negative binding examples (56.30%). We partitioned this data such that each cluster belonged to only one fold. A breakdown of each fold is shown in Table 1.

**Table 1:** Number of DTI examples and number of Morgan fingerprint clusters for each cross-validation fold and the test set used in the drug-based similarity split of the Human dataset.

| Partition | Total #<br>binding<br>examples | # positive<br>binding<br>examples | # negative<br>binding<br>examples | #<br>clusters |
| --- | --- | --- | --- | --- |
| CV Fold 0 | 1780 | 1267 | 513 | 1 |
| CV Fold 1 | 655 | 116 | 539 | 1 |
| CV Fold 2 | 1480 | 633 | 847 | 1 |
| CV Fold 3 | 861 | 84 | 777 | 1 |
| CV Fold 4 | 524 | 216 | 308 | 35 |
| Test set | 600 | 290 | 310 | 61 |

Despite the DrugBank drug-target interactions only being used for message passing and not for backpropagation, we sought to avoid leakage of binding information similar to that of the validation and test sets. We calculated the medioids of each of the 100 Morgan fingerprint clusters. We then generated Morgan fingerprints of all drugs in the DrugBank dataset and assigned each drug to the cluster whose medioid was the most similar per Tanimoto similarity. For each cross-validation fold, we only included DrugBank drug-target interactions as message passing edges if the drug was most similar to a cluster used in the fold’s training set.

### 2.5 Benchmarking

We selected four state-of-the-art, open-source deep learning models for DTI prediction against which to benchmark PIGLET: AMMVFDTI [35], FragXsiteDTI [13], TransformerCPI [5], and MSF-DTA [20]. AMMVF-DTI and TransformerCPI are both sequence-based DTI prediction models, FragXsiteDTI is a structure-based DTI prediction model, and MSF-DTA is a network-based DTI prediction model. We trained each model on the training set for the default number of epochs provided in the published code. We used AUROC on the validation set to select the best epoch for each iteration of each model. We computed the AUROC on the test set at the selected epoch only. For both of the split settings, we evaluated performance of each model using the average test AUROC for 20 independent runs (random split: 20 random seeds; drug split: 4 random seeds and 5 cross-validation folds).

### 2.6 Case study: 2025 FDA approvals

We sought to demonstrate utility of the PIGLET model on a case study exemplifying potential for real-world drug discovery. We identified 11 drugs that gained FDA approval in 2025 and had known targets in the canonical human proteome. As the DrugBank information used in constructing message-passing edges was from the 2023 version of DrugBank, none of these novel drugs had previously been seen in the graph. We assessed if PIGLET could identify the targets of any of these 11 novel drugs.

We froze the weights of the single best PIGLET model from cross-validation, as determined by validation AUROC. We used this model to calculate the drug-target interaction score between each of the 11 novel drugs and each of the 15,888 human proteins in the largest connected component of the proteome-wide interaction graph. We assessed if known targets were recovered by receiving high PIGLET scores.

## 3 Results

### 3.1 Proteome-wide interaction graph

Properties of the final proteome-wide interaction graph *G* are shown in Table 2.

**Table 2:** Properties of the proteome-wide interaction graph *G* and its largest connected component *H* .

| Property | Full graph $G$ | Largest connected component $H$ |
| --- | --- | --- |
| # nodes | 25,630 | 22,895 |
| # target nodes | 18,623 | 15,888 |
| # drug nodes | 7,007 | 7,007 |
| # edges | 3,604,984 | 3,604,206 |
| # target-drug binding edges (Human / prediction) | 5,900 | 5,900 |
| # target-drug binding edges (DrugBank / message passing) | 12,331 | 12,331 |
| # target-target PPI edges | 63,780 | 63,712 |
| # target-target similarity edges | 3,075,561 | 3,074,851 |
| # drug-drug similarity edges | 442,650 | 442,650 |
| Average degree | 281.31 | 314.85 |
| Average target node degree | 338.12 | 396.23 |
| Average drug node degree | 130.31 | 130.31 |
| # connected components | 2,200 | 1 |

### 3.2 All methods perform similarly well on random split

Consistent with previous work [8], we observed that all five methods evaluated had very strong performance on random splits of the Human dataset (Table 3). All methods had average test AUCs between 0.975 and 0.983. The gap between average validation AUC and average test AUC was small, ranging between 0.000 to 0.005.

**Table 3:** AUROCs on the validation and test sets of the Human dataset using random splits. Results are shown averaged over 20 random seeds, with standard deviation in parentheses.

| Model | Val AUROC | Test AUROC |
| --- | --- | --- |
| AMMVF-DTI | **0.983** (SD = 0.004) | 0.982 (SD = 0.004) |
| FragXsiteDTI | 0.979 (SD = 0.005) | 0.975 (SD = 0.006) |
| MSF-DTA | **0.983** (SD = 0.005) | **0.983** (SD = 0.005) |
| TransformerCPI | 0.971 (SD = 0.006) | 0.972 (SD = 0.007) |
| PIGLET (ours) | 0.980 (SD = 0.004) | 0.975 (SD = 0.006) |

### 3.3 PIGLET outperforms all other methods on drug similarity split

All methods showed a drop in performance when using the drug similarity split as opposed to the random split (Table 4). PIGLET performed the best, with an average test AUC of 0.873. MSF-DTA, the other network-based method, had the second-highest average test AUC (0.841). The three sequence- and structure-based methods had poor performance on the drug split, with mean validation AUCs ranging between 0.738 and 0.755, and mean test AUCs ranging between 0.531 and 0.642. All models showed larger variance in AUROC across runs than in the random split setting, likely driven by the cross-validation approach.

**Table 4:** AUROCs on the validation and test sets of the Human dataset using drug similarity splits. Results are shown averaged over five cross-validation folds and four random seeds, with standard deviation in parentheses.

| Model | Val AUROC | Test AUROC |
| --- | --- | --- |
| AMMVF-DTI | 0.755 (SD = 0.076) | 0.642 (SD = 0.148) |
| FragXsiteDTI | 0.738 (SD = 0.098) | 0.531 (SD = 0.056) |
| MSF-DTA | 0.850 (SD = 0.069) | 0.841 (SD = 0.045) |
| TransformerCPI | 0.740 (SD = 0.166) | 0.542 (SD = 0.113) |
| PIGLET (ours) | **0.928** (SD = 0.056) | **0.873** (SD = 0.019) |

### 3.4 Message-passing edges from DrugBank drive performance increases on drug similarity split

We examined differences in PIGLET performance when including and excluding the DrugBank message passing edges (Table 5). Performance on the random split was the same in both setups, suggesting that data leakage drives high performance and additional message-passing edges does not further improve the model. When using the drug similarity split of the Human dataset, a difference in AUROC between the graph setups with and without the DrugBank message passing edges emerged, with the former having a mean test AUC of 0.873 and the latter having a mean test AUC of 0.720.

**Table 5:** AUROCs on the validation and test sets of the Human dataset for both split settings, with and without the DrugBank message-passing edges. Results are shown averaged over 20 independent runs (20 random seeds for random split; 5 cross-validation folds and 4 random seeds for drug split), with standard deviation in parentheses.

| Model | Split | DrugBank MP? | Val AUROC | Test AUROC |
| --- | --- | --- | --- | --- |
| PIGLET | Random | Yes | 0.980 (SD = 0.004) | 0.975 (SD = 0.006) |
| PIGLET | Random | No | 0.981 (SD = 0.005) | 0.977 (SD = 0.006) |
| PIGLET | Drug | Yes | 0.928 (SD = 0.056) | 0.873 (SD = 0.019) |
| PIGLET | Drug | No | 0.858 (SD = 0.065) | 0.720 (SD = 0.115) |

### 3.5 PIGLET can recover targets of novel drugs

Out of the 21 known drug-target interactions annotated in DrugBank for the 11 drugs which gained FDA approval in 2025, six had a PIGLET score of 0.9 or higher (Table 6). Three of the 11 drugs had at least one ground truth target recovered with a score of 0.9 or higher.

**Table 6:** PIGLET scores for ground truth drug-target interactions for eleven drugs that gained FDA approval in 2025. The PIGLET score percentile indicates the fraction of targets with a higher PIGLET score than the given target.

| DrugBank ID | Drug Name | UniProt ID | Target Name | PIGLET Score | PIGLET Score Percentile |
| --- | --- | --- | --- | --- | --- |
| DB12134 | Gepotidacin | P11388 | DNA topoisomerase 2-alpha | 0.023278 | 0.638721 |
| DB12134 | Gepotidacin | Q02880 | DNA topoisomerase 2-beta | 0.033875 | 0.575843 |
| DB12580 | Tradipitant | P25103 | Substance-P receptor | 0.005112 | 0.421954 |
| DB13262 | Aceclidine | P08912 | Muscarinic acetylcholine receptor M5 | 0.000154 | 0.986405 |
| DB13262 | Aceclidine | P08173 | Muscarinic acetylcholine receptor M4 | 0.000563 | 0.963872 |
| DB13262 | Aceclidine | P20309 | Muscarinic acetylcholine receptor M3 | 0.005726 | 0.852027 |
| DB13262 | Aceclidine | P11229 | Muscarinic acetylcholine receptor M1 | 0.000192 | 0.982880 |
| DB13262 | Aceclidine | P08172 | Muscarinic acetylcholine receptor M2 | 0.000298 | 0.975579 |
| DB14844 | Dordaviprone | P35462 | D(3) dopamine receptor | 0.040114 | 0.958396 |
| DB14844 | Dordaviprone | P14416 | D(2) dopamine receptor | 0.001639 | 0.996916 |
| DB16133 | Delgocitinib | P23458 | Tyrosine-protein kinase JAK1 | 0.983630 | 0.372545 |
| DB16133 | Delgocitinib | O60674 | Tyrosine-protein kinase JAK2 | 0.958550 | 0.541100 |
| DB16133 | Delgocitinib | P52333 | Tyrosine-protein kinase JAK3 | 0.978544 | 0.423716 |
| DB16133 | Delgocitinib | P29597 | Non-receptor tyrosine-protein kinase TYK2 | 0.990501 | 0.276624 |
| DB16277 | Paltusotine | P30874 | Somatostatin receptor type 2 | 0.002738 | 0.928185 |
| DB17520 | Vimseltinib | P07333 | Macrophage colony-stimulating factor 1 receptor | 0.706939 | 0.261518 |
| DB18490 | Aficamten | Q14896 | Myosin-binding protein C, cardiac-type | 0.998205 | 0.509063 |
| DB18968 | Elinzanetant | P29371 | Neuromedin-K receptor | 0.000292 | 0.872545 |
| DB18968 | Elinzanetant | P25103 | Substance-P receptor | 0.005436 | 0.345984 |
| DB19043 | Imlunestrant | P03372 | Estrogen receptor | 0.955696 | 0.001070 |
| DB19202 | Acoltremon | Q7Z2W7 | Transient receptor potential cation channel subfamily M member 8 | 0.647182 | 0.568291 |

### 3.6 Network-based models are faster than sequence- and structure-based models

In addition to evaluating models based on classification performance, we also compared model runtimes. We calculated the average training time for each model over the 20 random split runs and 20 drug split runs. Here, training time is defined as the number of minutes until the model achieves the best validation AUROC; we did not include the time required to preprocess inputs. We show the average training times and standard error per model in Figure 2. PIGLET and MSF-DTA, both network-based methods, had the fastest training times. Both models completed training in less than 20 minutes on average. FragXSiteDTI had by far the slowest training time, completing training in about 4.8 hours on average.

**Figure 2:**
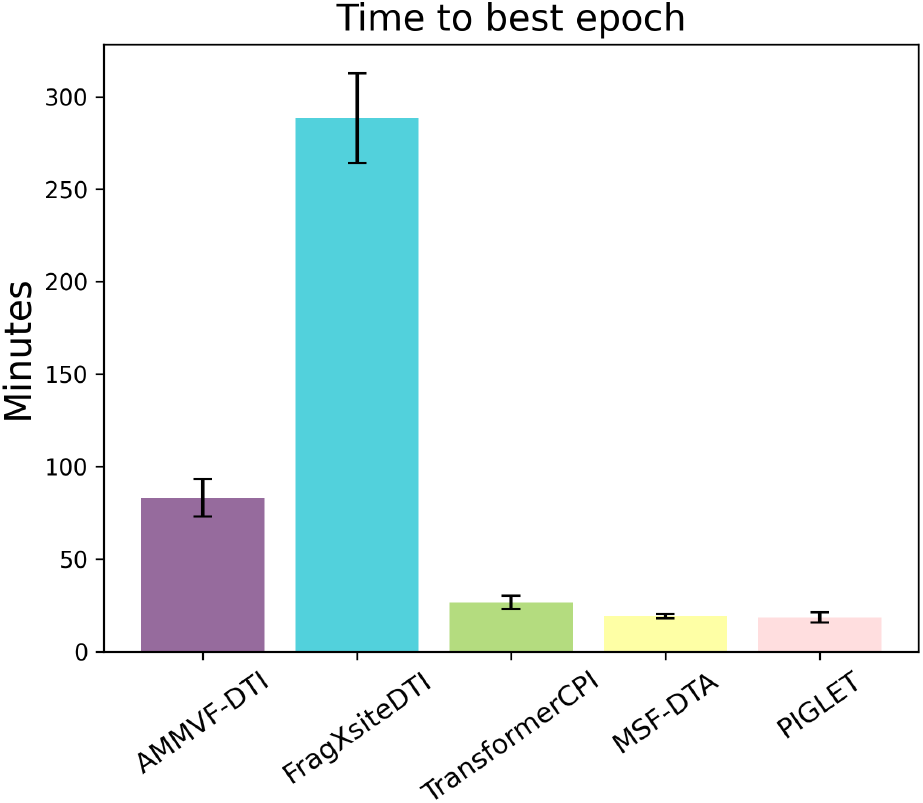
Average number of minutes to best epoch (per validation AUROC) for each model across a total of 40 runs. Standard error shown with error bars. The two network-based methods, PIGLET and MSF-DTA, have faster training time than the other three models.

## 4 Discussion

We have introduced PIGLET, a graph transformer method for predicting drug-target interactions. PIGLET leverages a knowledge graph of protein-protein interactions, target binding pocket similarity, and drug similarity in addition to drug-target binding relationships. We created a novel proteome-wide interaction graph using known and predicted binding pockets across the human proteome. We used information from DrugBank to guide the inductive bias of the proteome-wide interaction graph. We demonstrated that our GNN approach using this interaction graph outperforms existing deep learning methods for DTI on a drug similarity split of the Human dataset, and performs competitively with them on a random split. Additionally, we created a novel rigorous split of the Human dataset for evaluating DTI methods and presented an evaluation of existing DTI methods re-trained using the same data splits and training frameworks.

Many DTI prediction methods, including those which we used as benchmarks here, report extremely high performance using random splits of the Human dataset. Random splits often lead to data leakage and overestimation of model performance. In an attempt to ameliorate this issue, we created a split of the Human dataset based on drug embeddings. Beyond being more rigorous than a completely random interaction split, putting similar drugs in the same split is more challenging than randomly splitting by drug. As expected, models that previously reported very high performance on the random split had decreased performance on the drug-based split. We believe that using this split gives a more realistic evaluation of predictive methods.

While PIGLET and MSF-DTA were both top performers per test set AUC and runtime, and are both network-based DTI predictors, key differences between them suggest that PIGLET could have an advantage in predicting drug interactions with human proteins beyond those in the Human dataset. MSF-DTA leverages protein sequence similarity in its protein representations, PIGLET leverages local structural similarity. This distinction is important because highly similar protein structure motifs can be made by very different primary sequences. Additionally, comparing binding sites is preferable to comparing overall similarity because doing so accounts for the case where globally dissimilar proteins can bind the same ligand due to local binding site similarity.

Despite the relatively small size of the Human dataset, especially when reserving a validation set for epoch selection and when excluding the test set, our PIGLET model used for deployment was able to recover some ground truth target interactions of novel drugs. This result demonstrates preliminary progress toward real-world utility in drug-target interaction prediction beyond saturating performance on small, established benchmarks. However, our deployment model was limited by training data size and may perform better when trained on the full extent of drug-target interactions present in DrugBank and DTI datasets beyond the Human dataset.

While the DTI task is less granular and requires a choice of affinity threshold to define binary interaction labels, the DTA task is made challenging by inconsistencies and errors present in binding affinity data [12, 15, 34]. For this reason, we have chosen to approach the DTI task instead of the DTA task.

## 5 Limitations and Ethical Considerations

A limitation of this work is that we only benchmarked using the Human dataset. Other datasets, such as C. elegans [18], Davis [7], KIBA [27], and BindingDB [19], are commonly used to benchmark DTI models. We selected only the Human dataset due to our focus on drug interactions with human proteins for the purpose of early-stage drug development and drug repurposing. However, future work should repeat our standardized random and drug split evaluations of DTI methods using these additional datasets to more comprehensively compare methods against each other.

Similarly, we only benchmarked PIGLET against four existing DTI prediction models, despite the existence of many other models with excellent reported performance [8]. Because we sought to re-train existing models on our new data splits, we were limited to benchmarking against methods with code and reproducible dependencies available on GitHub. However, a broader comparison of re-trained open-source methods is warranted in future work.

Predictions from PIGLET should not be interpreted as medical facts. While we propose PIGLET as a tool for drug repurposing, hypothesized binding relationships from PIGLET obviously require additional computational and experimental validation before approaching the clinic. Beyond confirmation of a binding relationship, experiments to determine appropriate dosage, therapeutic functional effect, and safety in the proposed patient population are necessary.

## Acknowledgments

KAC is supported by NIH F31GM151783; RBA is supported by NIH R35GM153195 and Chan Zuckerberg Biohub. We thank Jure Leskovec, Gowri Nayar, Alp Tartici, Mohini Misra, Issah Samori, and Yao Xu for helpful discussion. We thank Yao Xu for assistance in procuring data necessary to retrain the MSF-DTA benchmark model. Most of the computing for this project was performed on the Sherlock cluster. We would like to thank Stanford University and the Stanford Research Computing Center for providing computational resources and support that contributed to these research results.

## GenAI Disclosure

No Generative AI was used in the writing of this manuscript. Generative AI was used to accelerate writing of code; all code and outputs were verified by an author before use for the results presented here.

